# Dissolved gases from pressure changes in the lungs elicit an immune response in human peripheral blood

**DOI:** 10.1101/2023.10.18.562856

**Authors:** Abigail G. Harrell, Stephen R. Thom, C. Wyatt Shields

**Affiliations:** Department of Chemical and Biological Engineering, University of Colorado Boulder, Boulder, CO 80303, United States; Department of Emergency Medicine, University of Maryland School of Medicine, Baltimore, MD 21201, United States; Biomedical Engineering Program, University of Colorado Boulder, Boulder CO 80303, United States

**Keywords:** Diving, Lung-on-a-chip, Organ-on-a-chip, Peripheral Blood Mononuclear Cells (PBMCs), Dissolved Gases, Hyperbaric Chamber

## Abstract

Conventional dogma suggests that decompression sickness (DCS) is caused by nitrogen bubble nucleation in the blood vessels and/or tissues; however, the abundance of bubbles does not correlate with DCS severity. Since immune cells respond to chemical and environmental cues, we hypothesized that the elevated partial pressures of dissolved gases drive aberrant immune cell phenotypes in the alveolar vasculature. To test this hypothesis, we measured immune responses within human lung-on–a-chip devices established with primary alveolar cells and microvascular cells. Devices were pressurized to 1.0 or 3.5 atm and surrounded by normal alveolar air or oxygen-reduced air. Phenotyping of neutrophils, monocytes, and dendritic cells as well as multiplexed ELISA revealed that immune responses occur within 1 hour and that normal alveolar air (i.e., hyperbaric oxygen and nitrogen) confer greater immune activation. This work strongly suggests innate immune cell reactions initiated at elevated partial pressures contribute to the etiology of DCS.

**Translational Impact Statement:** Our work reveals that the elevated partial pressures of dissolved gases within the vasculature drives immune cell activation. These findings have broad implications in identifying novel biomarkers for high-risk individuals (e.g., divers, pilots) through screening differential immune markers across patients. Further, this work offers a lens through which novel strategies to mitigate DCS in clinical scenarios may be employed by the delivery of prophylactic drugs that temper immune reactions.

## 1 Introduction

Decompression sickness (DCS) is thought to occur due to a rapid decrease in ambient pressure. Typically, symptoms are experienced upon ascending from underwater dives; however, other scenarios, such as high-altitude flying and spacecraft extravehicular activity, can also cause DCS.^1,2^ While rarely fatal, symptoms include headache, visual changes, rash, dyspnea, and joint pain that can risk the safety of individuals.^3^ In the context of diving, the partial pressures of nitrogen and oxygen increase in the lungs, which results in elevated gas concentrations in the blood. During ascension, nitrogen bubbles can nucleate from gas supersaturation, which is thought to cause the symptoms related to DCS.^4^ While the impacts of these bubbles depend upon the organ system, the primary impacts are mechanical tissue damage, tissue compression, and obstruction of blood vessels, which have been shown to influence cellular activation and biochemical pathways.^5–7^

In some cases, DCS is monitored by precordial ultrasound where venous gas emboli (VGE) are detected.^8^ However, while DCS and VGE correlation has been identified, DCS symptoms have been reported in the absence of VGE.^9^ This lack of specificity obfuscates diagnostic efforts and the potential etiology of DCS.^10^ Additionally, individual variance in responses are often reported, exacerbating efforts to predict and diagnose DCS. DCS is traditionally viewed as related to gas bubble formation from insoluble gas on decompression.^11,12^ The inconsistent presence of bubbles in human studies has prompted investigations focused on inflammatory pathways in the blood.^13–15^ A large body of work implicates a subset of extracellular vesicles (EVs), 0.1 to 1.0 μm microparticles (MPs), that are elevated in humans and rodents exposed to high gas pressures and rise further after decompression.^7,16–26^ Inflammatory MPs production is an oxidative stress response triggered by gases associated with diving in a potency series of Argon (Ar)> Nitrogen (N_2_) >> Helium (He).^6^

While both *in vitro* and *in vivo* models have been used to assess immune responses to pressure changes and nucleated bubbles, they fail to account for the role of dissolved gases in a physiologically relevant manner.^27–29^ Thus, we report a model that recapitulates the physiology of the human lungs during compression and decompression using human lung-on-a-chip devices. These devices mimic gas permeation through the alveolar epithelium due to elevated hydrostatic pressures, gas dissolution in whole blood, and interactions between circulating blood and the vascular endothelium (Table 1). Along with the human lung-on-a-chip devices, we recapitulated the physiological changes that occur during underwater diving by altering the gas composition surrounding the human lung-on-a-chip devices to reflect the partial pressures of nitrogen, oxygen, and carbon dioxide within the lungs at a dive depth of 25 m. Innate immune cells are the first line of defense and protect the body from a variety of invaders and conditions, and they are known to respond to chemical and physical cues in minutes to hours.^30,31^ Therefore, we screened three populations of innate immune cells in blood: neutrophils, monocytes, and dendritic cells. By combining (i) whole blood perfusion in the human lung-on-a-chip devices, (ii) gas composition adjustments to match alveolar air at the prescribed dive depths, (iii) a hyperbaric chamber, and (iv) multiple phenotyping tools (i.e., flow cytometry and multiplexed ELISA), significant immune activation was observed.

**Table 1.**
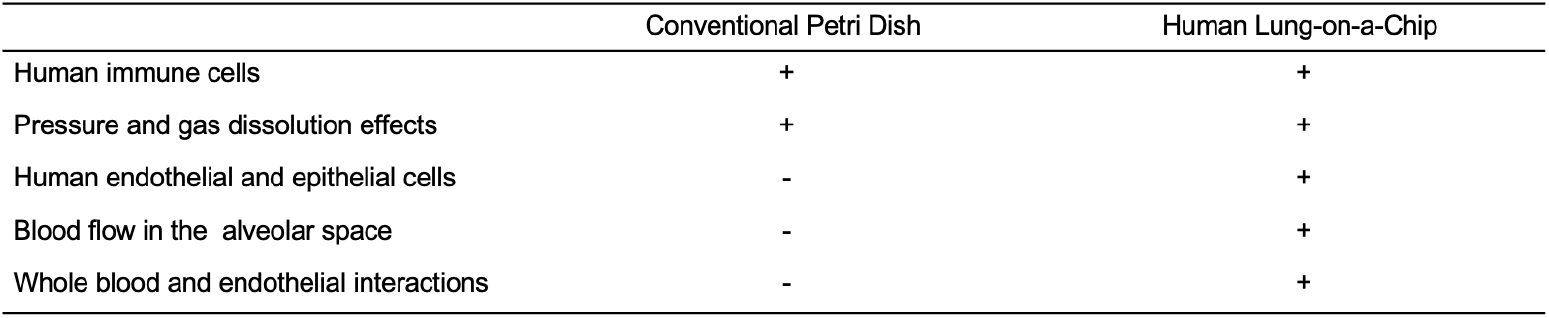
Traditional petri dish and human lung-on-a-chip device comparison.

|  | Conventional Petri Dish | Human Lung-on-a-Chip |
| --- | --- | --- |
| Human immune cells | + | + |
| Pressure and gas dissolution effects | + | + |
| Human endothelial and epithelial cells | - | + |
| Blood flow in the alveolar space | - | + |
| Whole blood and endothelial interactions | - | + |

## 2 Results and Discussion

### 2.1 Establishment and design of a physiological system

We demonstrate an *in vitro* experimental system involving human lung-on-a-chip devices that can mimic the physiology of the human lungs during diving to determine the role of gas dissolution and pressure effects on innate immune responses (Figure 1A). These devices have two channels made from polydimethylsiloxane (PDMS) with a porous membrane separating the top and bottom channels. This membrane permits gas exchange which is crucial for our system since gas dissolution is hypothesized to play a role in decompression sickness (DCS) pathophysiology (Figure 1B).

**Figure 1.**
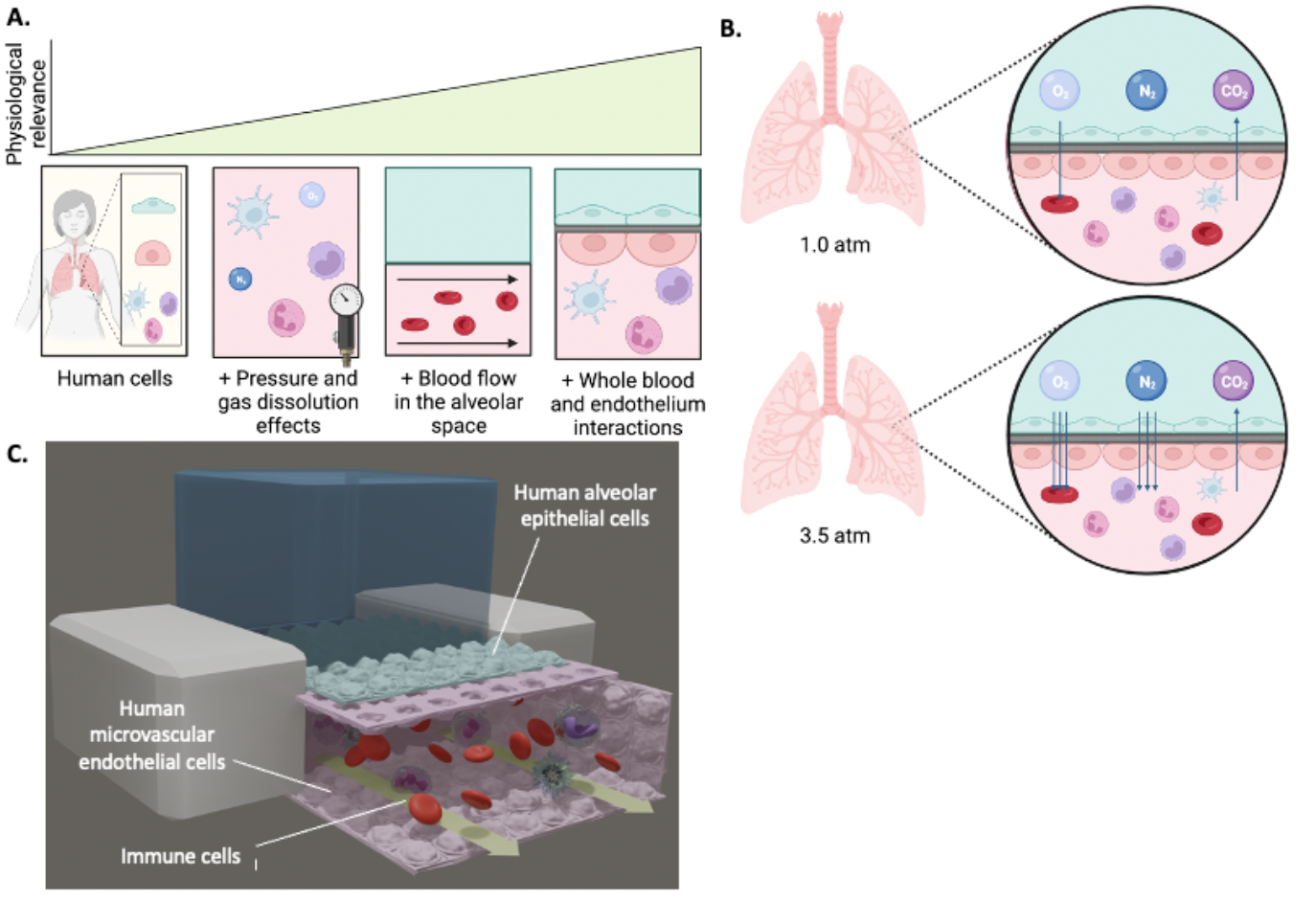
Microchip model of the human lung to study immune responses to changes in gas dissolution from elevated pressure. (A) Decompression sickness is driven by changes in partial pressures of gases in blood leading to increased dissolved O_2_ and N_2_. (B) Human lung-on-a-chip devices recapitulate interactions between whole blood and the vascular endothelium, effects from gas dissolution through the epithelium due to pressure fluctuations, and blood flow in the alveolar space. (C) Cross-sectional schematic illustration of a human lung-on-a-chip device.

After establishing the lung-on-a-chip devices (Figure 1C), we infused human peripheral blood into the bottom channel of the chip and exposed the bottom channel to the appropriate shear stresses experienced in the vasculature of alveoli (Figure 2A and B). We designed a hyperbaric chamber that can pressurize our human lung-on-a-chip devices for at least one hour. This chamber was outfitted with a needle valve to allow for a controlled rate of decompression at 0.9 atm/minute as recommended by diving protocols.^32^ We chose to follow this countermeasure to focus on the role of gas dissolution and pressure effects rather than the immunogenicity of bubbles that form during decompression. Furthermore, since the partial pressure of gases in the alveolar space is dependent upon the ambient pressure that changes during dives, we chose to surround our samples with corresponding gas compositions in the lungs at either 1.0 or 3.5 atm based on the experimental pressure condition (Figure 2C). An additional “oxygen-reduced” gas composition group was studied to disentangle the roles of hyperbaric nitrogen and oxygen toxicity. In this group, the partial pressure of oxygen was reduced to a level where its partial pressure at 3.5 atm matches the partial pressure of oxygen at 1.0 atm (Table 2). Three innate peripheral blood mononuclear cell (PBMC) populations were chosen for analysis (i.e., neutrophils, monocytes, and dendritic cells) because these cells can adapt their phenotypes over short timescales (Figure 2D). Also, we measured the secretion of 14 cytokines and chemokines to identify biochemical signals involved in the pathophysiology of DCS (Figure 2D). We compared the phenotyping results from our human lung-on-a-chip devices to a traditional 2D *in vitro* model by exposing human whole blood in a petri dish to the experimental conditions shown in Table 1. In comparison to the lung-on-a-chip data, data from the petri dish model (i) shows the importance of using a physiologically relevant model and (ii) decouples some of the variance that is introduced from blood flow interacting with endothelial cells.

**Table 2.** Alveolar Gas Mixtures. Assumes that only N_2_, O_2_, and CO_2_ are present.

| **Pressure** |  | **1 atm** | **3.5 atm** | **3.5 atm** |
| --- | --- | --- | --- | --- |
|  |  | **Normal Alveolar Air** | **Normal Alveolar Air** | **Oxygen-Reduced Air** |
| **(Dive Depth)** |  | **(0.0 m)** | **(25.1 m)** | **(25.1 m)** |
| **Partial pressures (mmHg)** | N<sub>2</sub> | 573.0 | 2005.5 | 2005.5 |
|  | O<sub>2</sub> | 100.0 | 350.0 | 100.0 |
|  | CO<sub>2</sub> | 40.0 | 40.0 | 40.0 |
| **Gas composition (percent)** | N<sub>2</sub> | 80.4% | 83.7% | 93.5% |
|  | O<sub>2</sub> | 14.0% | 14.6% | 4.7% |
|  | CO<sub>2</sub> | 5.6% | 1.7% | 1.9% |

**Figure 2.**
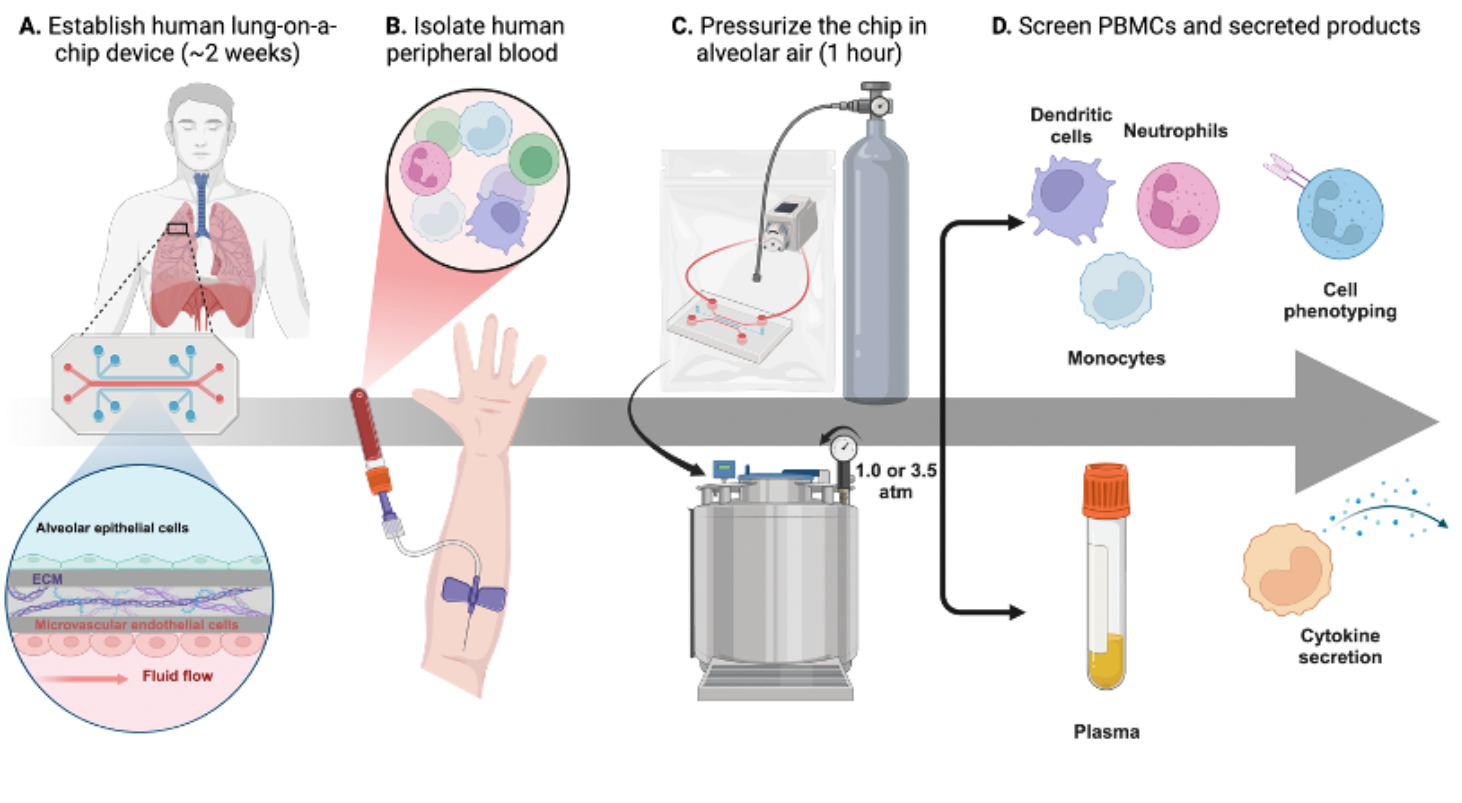
Establishment, pressurization, and analysis of human lung-on-a-chip devices. (A) Graphical illustration of establishing the human lung-on-a-chip devices. (B) Blood is collected from healthy human volunteers and infused into the bottom channel of the device. (C) The bottom channel is connected to a battery-operated peristaltic pump, placed into a gas-impermeable bag, and flushed with the appropriate gas mixture. The bag is placed into the hyperbaric chamber and pressurized. (D) After 1 hour, the chamber is vented at 0.9 atm/min, and plasma, dendritic cells, monocytes, and neutrophils are extracted, processed, and analyzed using multiplexed ELISA and flow cytometry.

### 2.2 Neutrophil phenotyping shows significant changes in proinflammatory and anti-inflammatory markers when exposed to elevated pressures

Both proinflammatory and anti-inflammatory markers were assessed to determine neutrophil activation (Figure 3A). For the lung-on-a-chip devices, there was significant upregulation of most neutrophil markers including NE, MPO, and CD41a (Figure 3B); CD18 displayed only minor changes. Both NE and MPO are proinflammatory markers with roles in mucus secretion and reactive oxygen and nitrogen species production, respectively.^33,34^ However, CD41a is an anti-inflammatory marker that indicates platelet-neutrophil attachment, thus suggesting platelet activation and cell adhesion.^35^ The upregulation of both anti-inflammatory and proinflammatory markers indicates that this response is more complex than the traditional N1/N2 dichotomy. There were few significant differences observed between the oxygen-reduced and normal alveolar air conditions at 3.5 atm, indicating that responses are predominantly reliant upon elevated nitrogen partial pressures rather than oxygen toxicity. Out of the three cells that were phenotyped, neutrophil markers showed the most significant changes between conditions; therefore, neutrophils may be the best choice for future diagnostic assays. Specifically, NE and CD41a could be used as future diagnostic markers as there is a clear and significant upregulation for both in the 3.5 atm normal and 3.5 oxygen-reduced alveolar air conditions. Furthermore, considering the existing research on neutrophil microparticle formation due to elevated pressures, further testing of neutrophil markers may be a good complement to these studies.^7^

**Figure 3.**
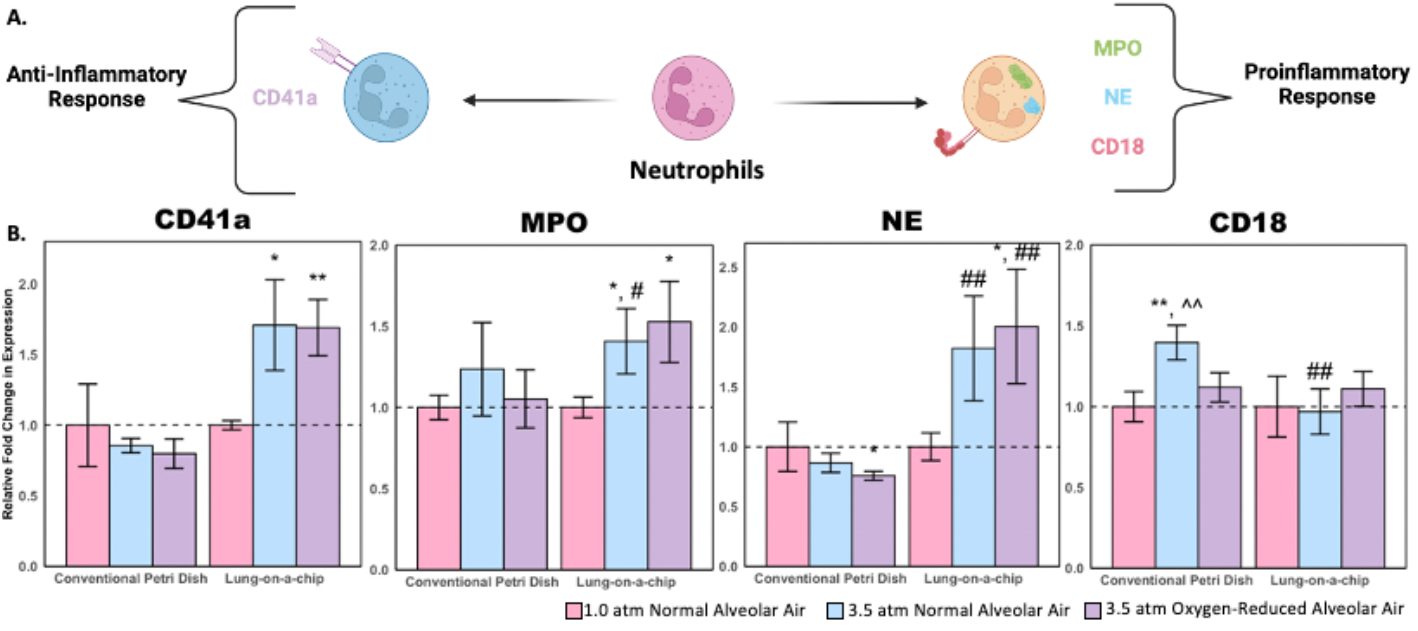
Neutrophil phenotyping results. (A) Schematic illustration of neutrophil polarization showing proinflammatory and anti-inflammatory markers. (B) Relative expression of neutrophil activation markers normalized to 1.0 atm controls (SE n = 5; * P < 0.1 with respect to 1.0 atm controls; ** P < 0.05 with respect to 1.0 atm controls; # P < 0.1 with respect to the corresponding petri dish condition; ## P < 0.05 with respect to the corresponding petri dish condition; ^^ P < 0.05 with respect to the 3.5 atm oxygen-reduced condition).

### 2.3 Monocyte phenotyping shows that physiologically relevant models more accurately recapitulate the immune response to dissolved gases

A panel of both proinflammatory and anti-inflammatory markers were also analyzed for monocytes (Figure 4A). Of the four monocyte markers studied, arginase-1 (Arg-1), an anti-inflammatory marker, showed a three-fold change in expression for the 3.5 atm normal alveolar air condition and a nearly two-fold change in expression for the 3.5 atm oxygen-reduced alveolar air condition (Figure 4B). However, there were no significant changes in Arg-1 for any of the petri dish conditions studied. Despite the lower variability for the petri dish experiments, the significant upregulation of Arg-1 in the human lung-on-a-chip device studies shows that physiological conditions are crucial in developing immune cell responses to pressure and gas dissolution effects, which are not readily observed in a 2D *in vitro* model. This is further supported by the significant differences between petri dish and lung-on-a-chip analogs for both the oxygen-reduced and normal alveolar air conditions in the expression of CD80. Interestingly, HIF-1*α*showed similar trends across the two models, yet only showed significant differences in the petri dish condition. In this case, the petri dish experiments highlight potential trends that were not significant for the lung-on-a-chip models due to variability introduced from a more complex model. This marker indicates that petri dish experiments can be used to identify markers that are not necessarily reliant on physiologically relevant surroundings. HLA DR/DP, HIF-1*α*, and CD80 expression highlighted significant differences between the 3.5 atm normal alveolar air and 3.5 atm oxygen-reduced alveolar air in the petri dish models; however, further studies may be needed to study the variability seen in the lung-on-a-chip models to better identify differences between oxygen toxicity and increased nitrogen partial pressure effects in monocyte markers.

**Figure 4.**
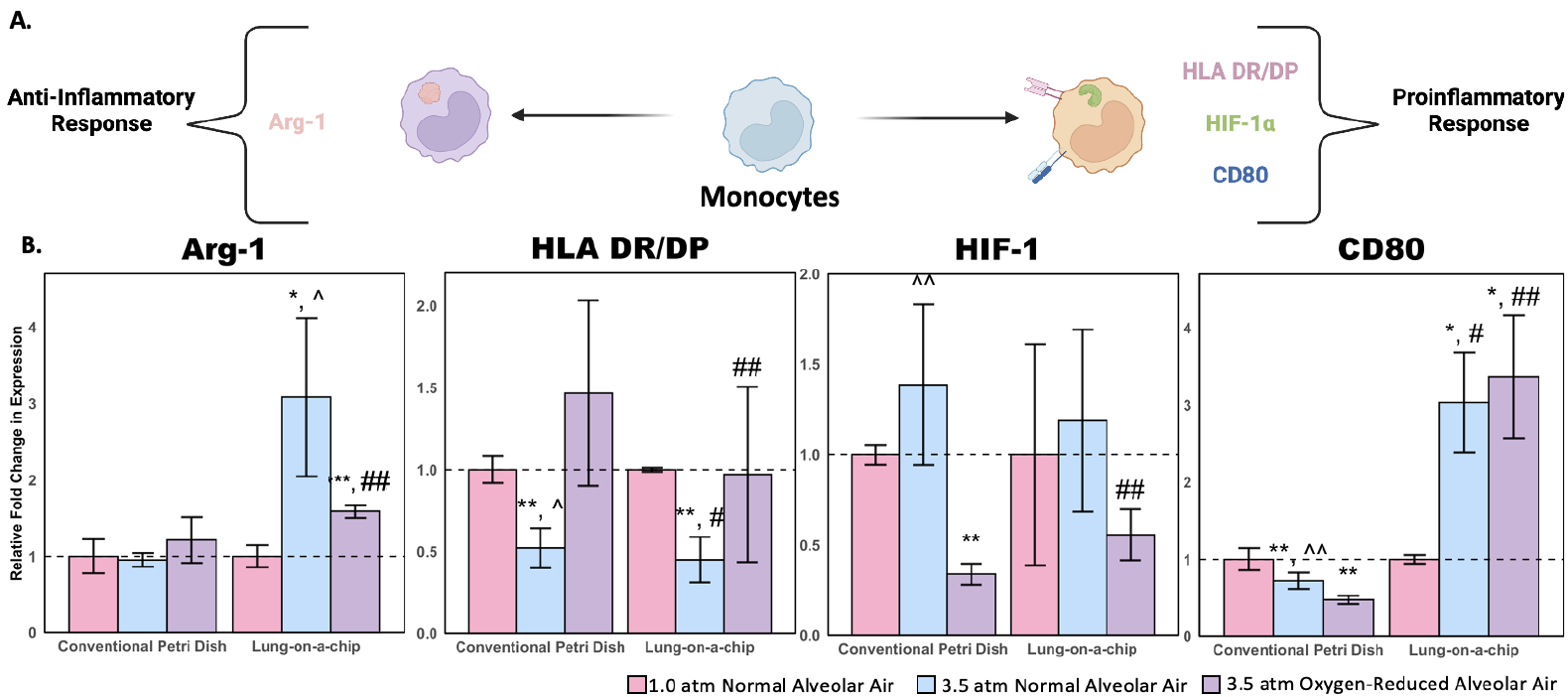
Monocyte phenotyping results. (A) Schematic illustration of monocyte polarization showing proinflammatory and anti-inflammatory markers. (B) Relative expression of monocyte activation markers normalized to 1.0 atm controls (SE n = 5; * P < 0.1 with respect to 1.0 atm controls; ** P < 0.05 with respect to 1.0 atm controls; # P < 0.1 with respect to the corresponding petri dish condition; ## P < 0.05 with respect to the corresponding petri dish condition; ^ P < 0.1 with respect to the 3.5 atm oxygen-reduced condition; ^^ P < 0.05 with respect to the 3.5 atm oxygen-reduced condition).

### 2.4 Dendritic cell phenotyping shows that earlier responders are more likely to show significant changes to dissolved gases

Similar to the neutrophil and monocyte phenotyping, multiple dendritic cell markers were assessed (Figure 5A). Dendritic cells showed the most variability across cell types for both the petri dish and lung-on-a-chip models but showed few differences between the oxygen-reduced and normal alveolar air conditions (Figure 5B). We measured the expression of HLA DR/DP, a protein receptor associated with antigen presentation, as well as CD80 and CD86, both of which are costimulatory molecules that play a crucial role in guiding adaptive immune responses. The lack of significant upregulation of these molecules suggest that they may be less sensitive to changes in gas partial pressures or that the 1 hour timescale of these experiments was insufficient to observe dendritic cell activation. Dendritic cells are the slowest-acting of the PBMC types studied, which corroborates findings from both the petri dish and lung-on-a-chip models.^36^ Therefore, longer time points or increased hydrostatic pressures may be needed to better understand how dendritic cells respond to elevated gas pressures. However, early trends with CD80 in the human lung-on-a-chip devices suggest that dendritic cells may become activated over longer exposures to elevated gas pressures, especially in oxygen-reduced conditions.

**Figure 5.**
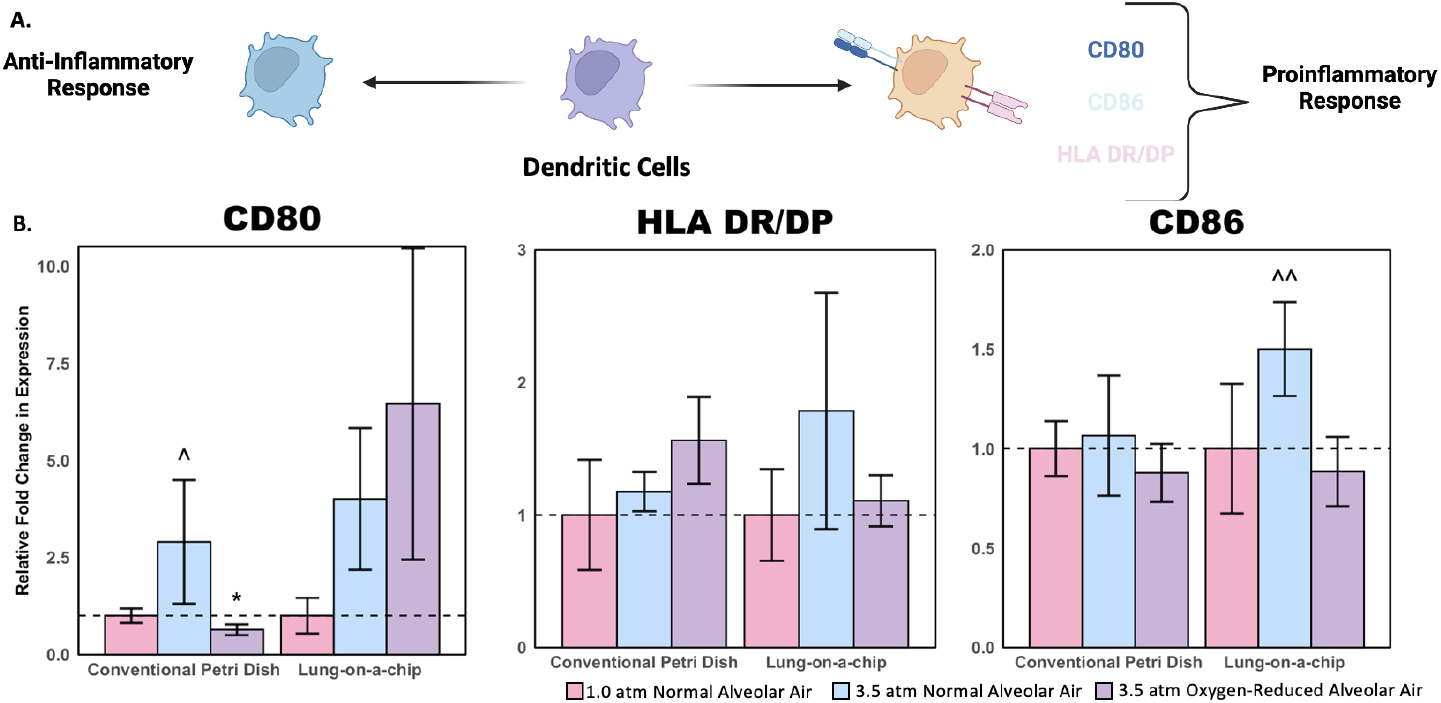
Dendritic cell phenotyping results. (A) Schematic illustration of dendritic cell polarization showing proinflammatory and anti-inflammatory markers. (B) Relative expression of dendritic cell activation markers normalized to 1.0 atm controls (SE n = 5; * P < 0.1 with respect to 1.0 atm controls; # P < 0.1 with respect to the corresponding lung-on-a-chip condition; ^ P < 0.1 with respect to the 3.5 atm oxygen-reduced condition; ^^ P < 0.05 with respect to the 3.5 atm oxygen-reduced condition).

### 2.5 Cytokine secretion data shows significant differences between elevated nitrogen partial pressure and oxygen toxicity impacts

We screened a variety of cytokines to identify potential diagnostic markers for an immune response during pressurization (Figure 6). Similar to the flow cytometry data, there was a mix of proinflammatory and anti-inflammatory responses, and few of the changes were significantly different. MIP-1*α* and MIP-1β were downregulated in the lung-on-a-chip models compared to 1.0 atm controls, suggesting that leukocyte recruitment to the endothelium decreased. Since cytokine secretion can occur over several hours, it is possible that more cytokines are being produced, but not yet secreted, in the timeline of these experiments. Generally, there was more upregulation of cytokines and chemokines for normal alveolar air at 3.5 atm in the lung-on-a-chip devices (showing the impacts of oxygen toxicity) or downregulation for oxygen-reduced alveolar air at 3.5 atm (showing the impacts of nitrogen partial pressures) with a few significant examples including IFN*α*, GM-CSF, and IL-1β. The petri dish conditions displayed more significant results potentially showing that the lung-on-a-chip devices introduced more variability, similar to studies on whole humans and murine models.^9^ This may be the result of endothelial cell production of cytokines or a response due to leukocyte-endothelial interactions, further emphasizing the importance of using a physiologically relevant model. Additionally, these data more clearly identify differences between oxygen-reduced alveolar air and normal alveolar air conditions. Across conditions, there is more cytokine production for the normal alveolar air (which is associated with oxygen toxicity) than oxygen-reduced air at 3.5 atm, which further contributes to the established perspective that hyperoxic conditions contribute to increased cytokine production and signal for additional immune cell mobilization.^37,38^ Furthermore, these data shows that endothelial cells and other physiological conditions are influencing the immune cell responses in ways that are not immediately identified from the flow cytometry data alone and support the need for a physiological model that replicates intravascular conditions.

**Figure 6.**
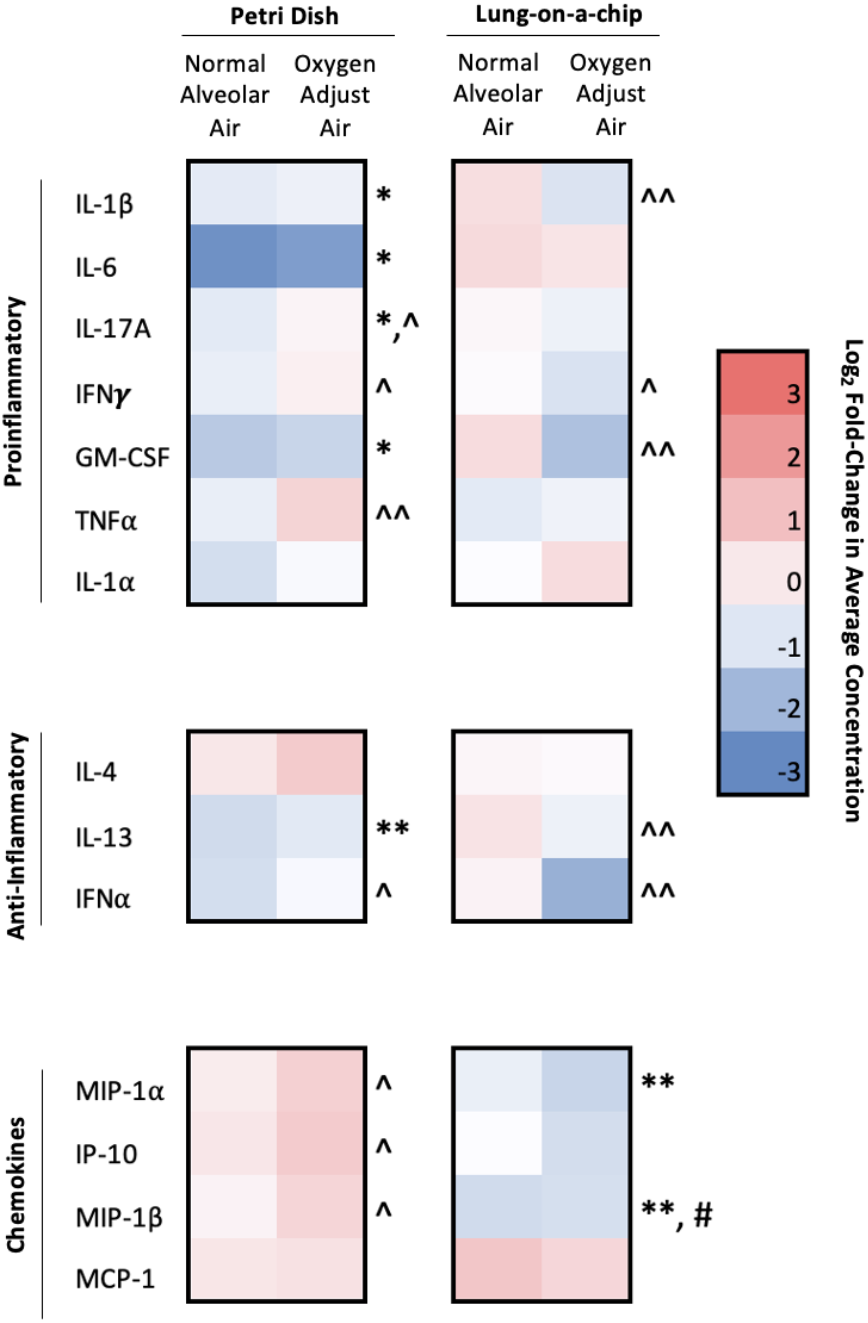
Cytokine secretion results. Cytokine secretion from cells in human lung-on-a chip devices and blood-containing petri dish models (* P < 0.1 with respect to 1.0 atm controls; ** P < 0.05 with respect to 1.0 atm controls; # P < 0.1 with respect to the corresponding petri dish condition; ^ P < 0.1 with respect to the 3.5 oxygen-reduced condition; ^^ P < 0.05 with respect to the 3.5 oxygen-reduced condition).

## 3 Discussion

Human lung-on-a-chip devices revealed significant immune activation in response to elevated gas partial pressures. Our experiments occurred over 1 hour, and cellular activity was immediately arrested after decompression using a metabolic inhibitor (N-ethylmaleimide). Since the synthesis and secretion of cytokines and cell expression of new protein receptors on the membrane are time-dependent, our results indicate that immune responses occur prior to decompression. These events play a critical role in the onset and etiology of DCS. From the flow cytometry data, dissolved gases can activate a variety of innate immune cells while the cytokine secretion data showed distinct differences between effects from increased oxygen vs. nitrogen partial pressures. Further, these data illustrate the importance of using a physiologically relevant model compared to traditional 2D models, as significant differences between the two models were found across our analyses. Future work may introduce longer timepoints to allow for cells to reach a steady state of secretion and activation. Additionally, epigenomics and transcriptomics should be performed on cells in these devices to determine the pathways of gene expression that mediates the cellular responses observed.

## 4 Materials and Methods

### 4.1 Hyperbaric system development

A hyperbaric chamber was built from custom parts including a compressor (California Air Tools) connected to an American Society of Mechanical Engineers (ASME)-code pressure tank (McMaster-Carr). The tank is certified to hold 7.5 atm, and it was outfitted with a needle-valve regulator (McMaster-Carr) to control the rate of gas expulsion for venting (i.e., at a controlled rate of 0.9 atm/min, which corresponds to the maximum allowed rate of ascension for divers).^32^

### 4.2 Cell culture

Primary human alveolar epithelial cells (HPAECs, CellBiologics) were cultured at 37°C and 5% CO_2_ with SABM medium (Lonza), SAGM supplements (Lonza), and 5% FBS (Thermo Fisher) in a T-25 flasks coated with a gelatin solution (Sigma-Aldrich) until confluent. Primary human lung microvascular endothelial cells (HMVEC-Ls, Lonza) were cultured at 37°C and 5% CO_2_ with EBM-2 medium (Lonza) and EGM-2MV supplements in T-75 flasks until confluent. As recommended by both manufacturers, no cell line was used beyond Passage 5.

### 4.3 Lung-on-a-chip device establishment

Chip establishment was performed according to the instructions from the manufacturer (Alveolus Lung-Chip Co-Culture Protocol, Emulate). Briefly, each channel was flushed with ER-1 solution (Emulate), and the chips were incubated in a UV-light box to functionalize the top and bottom channel to allow for the extracellular matrix components (ECM) components to bind. The top channel was coated with an ECM mixture in DPBS with 200 μg/mL collagen IV (Sigma), 30 μg/mL fibronectin (Corning), and 5 μg/mL laminin (Sigma). The bottom channel was coated with an ECM mixture in DPBS with 200 μg/mL collagen IV and 30 μg/mL fibronectin. Chips incubated overnight at 37°C and 5% CO_2_. Each channel was washed with SAGM culture medium prior to cell seeding. Primary human alveolar epithelial cells (CellBiologics) were seeded into the top channel at a concentration of 1 x 10^6^ cells/mL and incubated overnight. Two days later, primary human lung microvascular endothelial cells (Lonza) were seeded in the bottom channel at a concentration of 5 x 10^6^ cells/mL. After one day of incubation, the bottom channel was connected to a peristaltic pump (NexTage) with 0.5 mm x 0.8 mm silicone tubing (NexTage) and exposed to constant media flow at 30 μL/hr. On Day 5 of culture, an air-liquid interface was introduced by aspirating the media from the top channel and replacing the bottom channel media with Media 199 (Thermo Fisher) supplemented with 10 ng/mL of human epidermal growth factor (Corning), 3 ng/mL human basic fibroblast growth factor (Corning), 0.135 ng/mL human vascular endothelial growth factor (Sigma), 1 μg/mL hydrocortisone (Sigma), 10 μg/mL heparin (Sigma), 80 μM dibutyryl cAMP (Sigma), 1 mM L-glutamax (Thermo Fisher), 20 nM dexamethasone (Sigma), 1% penicillin-streptomycin (Sigma), and 2% FBS (Sigma). Chips were ready to use after seven days of culture.

### 4.4 Lung-on-a-chip pressurization

Whole blood was obtained from healthy human donors by venipuncture and collected in BD Vacutainer™ plastic blood collection tubes with K2 EDTA: Hemogard™ Closure (Thermo Fisher) to prevent coagulation following an approved protocol from the IRB at the University of Colorado Boulder (22-0175). Chips were infused with 50 μL of the collected blood. The bottom channel was connected to a battery-operated peristaltic pump and flowed at a rate of 30 μL/hr. The chip and pump were placed in a Kynar gas-impermeable bag (Cole-Parmer) and flushed with the appropriate gas mixture (Table 2). The bag was placed into the pressure canister and pressurized to either 1.0 or 3.5 atm, as ambient pressure in Boulder, CO is 0.8 atm. After 1 hour under pressure, the chamber was depressurized at a rate of 0.9 atm/min. Whole blood was extracted from the bottom channel, and the bottom channel was flushed with 1 mL of DPS. The mixture of extracted blood was mixed with 5 μL of 10 mM N-ethylmaleimide (ThermoFisher) in DPBS following previous methods for arresting cellular metabolic activity.^39^

### 4.5 Petri dish pressurization

Whole blood was collected from healthy human donors in the same manner as above, and 1 mL of blood was seeded into a well of a 12-well plate. The plate was placed in a Kynar gas-impermeable bag and flushed with the appropriate gas mixture (Table 2). The bag was placed into the pressure tank and pressurized to either 1.0 or 3.5 atm. After 1 hour, the chamber was depressurized at a rate of 0.9 atm/min. Whole blood was extracted from the plate, and the plate was washed with 1 mL of DPS and added to the extracted whole blood. The mixture of extracted blood was mixed with 5 μL of 10 mM N-ethylmaleimide.

### 4.6 Peripheral blood mononuclear cell and neutrophil isolation

For the human lung-on-a-chip experiments, peripheral blood mononuclear cells (PBMCs) were isolated with EasySep™ Release Human CD45 Positive Selection Kit (StemCell), and neutrophils were isolated with EasySep™ Direct Human Neutrophil Isolation Kit (StemCell) according to the instructions from the manufacturer. For the petri dish experiments, PBMCs were isolated with Ficoll-Paque (Thermo Fisher), a density gradient medium, according to the instructions from the manufacturer.

### 4.7 Cellular phenotyping

Isolated cells were collected in 1.5 mL Eppendorf tubes, centrifuged (N.B., centrifugation steps in this study were all performed at 350xG for 5 min at 4°C unless otherwise noted), and resuspended in stain buffer (2% FBS in DPBS). Cells were centrifuged again, resuspended in 100 μL of human Fc Block (BD), and incubated for 30 min at 4°C. Next, 0.5 mL of stain buffer was added to wash the cells and pelleted by centrifugation. The pellet was resuspended in 1 mL stain buffer and separated into five new Eppendorf tubes.

Three of the Eppendorf tubes were centrifuged and resuspended in 1 mL PBS. The tubes were centrifuged again and resuspended in 100 μL Zombie Green viability stain solution (BioLegend) at room temperature, in the dark, for 30 min. Next, 0.5 mL of PBS was added to wash the cells to wash the cells, and tubes centrifuged again. After aspiration, 1 mL of stain buffer was added to resuspend the cells, and the tubes were centrifuged again to pellet cells. The cells were resuspended in 100 μL of extracellular stain solution, in the dark, for 30 min at 4°C. A description of each antibody used is included in Table S1 of the Supporting Information. The amount of each antibody used followed instructions from the manufacturer. Then, 0.5 mL of PBS was added to wash the cells, the tubes were centrifuged, and 1 mL of stain buffer was added, and the tubes were pelleted by centrifugation. Cells were resuspended in 400 μL of stain buffer and stored on ice until analysis by flow cytometry.

Two of the Eppendorf tubes were centrifuged and cells were resuspended in 250 μL of Fixation and Permeabilization (BD) solution for 20 min at room temperature. Cells were diluted with 0.5 mL 1x Perm/Wash™ (BD) and were pelleted by centrifugation. Then, 1 mL Perm/Wash™ solution was added, and the tubes centrifuged again to pellet cells. The cells were resuspended in 100 μL intracellular stain solution at room temperature, in the dark, for 30 min. A description of each antibody used is included in Table S1 of the Supporting Information. The amount of each antibody used followed instructions from the manufacturer. Then, 0.5 mL Perm/Wash™ solution was added to wash the cells, the tubes were centrifuged, 1 mL of stain buffer was added, and the cells were pelleted again by centrifugation. Cells were resuspended in 400 μL stain buffer and stored on ice until analysis by flow cytometry.

### 4.8 Multiplexed ELISA

Blood plasma extracted from each group (n=5) was analyzed using the Inflammation 20-Plex Human ProcartaPlex™ Panel (Invitrogen). Briefly, 25 μL of each sample was incubated with the bead mix and washed multiple times using a magnetic handheld plate washer. Then, 25 μL of 1X detection antibody mixture was added to each sample and incubated for 30 min Each sample was diluted with 50 μL of streptavidin phycoerythrin and incubated for 30 min. Samples were analyzed with a MAGPIX instrument (Luminex).

### 4.9 Statistical analysis

Data in Figures 3-6 are shown as mean fold-change ± standard error (SE). For determination of statistical significance, one tailed, unpaired t-tests were used. Significance was determined at the cutoff point * P < 0.1 and ** P < 0.5 given the short duration (1 hour) of exposing cells to elevated pressures.

## Acknowledgements

The authors thank Dr. Joseph Dragavon in the Light Microscopy core for his assistance in imaging the human lung-on-a-chip devices. This work was primarily supported by the Office of Naval Resource (ONR) Undersea Medicine Program (N000142212541). Additional support by the NIH (R35GM147455, R21CA267608) and the National Science Foundation (NSF CBET 2143419) is acknowledged. C.W.S. is a Pew Scholar in the Biomedical Sciences, supported by the Pew Charitable Trusts. C.W.S. would also like to thank the Packard Foundation for their support of this project. The authors acknowledge support from the BioFrontiers Institute Advanced Light Microscopy Core (RRID: SCR_018302) and the Flow Cytometry Shared Core (S10ODO21601) at the University of Colorado Boulder. Some of the figures in this article were made using BioRender.com. The authors declare no conflict of interest.

