## Supporting Information for "Dissolved gases from pressure changes in the lungs elicit an immune response in human peripheral blood"

Supplementary Materials

Table S1. Antibodies used for flow cytometry.

Supplementary Figures

Figure S1. Panel organization for distinguishing and phenotyping monocytes, dendritic cells, and neutrophils.

Figure S2. Immune cell gating.

Figure S3. Standard curves from multiplexed ELISA.

### I. Supplementary Materials

**Table S1. Antibodies used for flow cytometry.** Extracellular antibodies are indicated with an asterisk (*) and intracellular antibodies are represented with a hashtag ($^{\#}$).

| Antibody target | Fluorophore | Host/isotype | Clone | Supplier |
|---|---|---|---|---|
| CD11b* | PE-Cyanine7 | Mouse / IgG1, kappa | ICRF44 | ThermoFisher |
| CD11c* | PE-Cyanine5.5 | Mouse / IgG1, kappa | 3.9 | ThermoFisher |
| CD80* | Super Bright 436 | Mouse / IgG1, kappa | 2D10.4 | ThermoFisher |
| HLA DR/DP* | Super Bright 600 | Mouse / IgG2a | HL-38 | ThermoFisher |
| CD86* | PE | Mouse / IgG2b, kappa | IT2.2 | ThermoFisher |
| CD14* | PE-Cyanine5 | Mouse / IgG1, kappa | 61D3 | ThermoFisher |
| Arg-1$^{\#}$ | eFluor 450 | Rat / IgG2a, kappa | A1exF5 | ThermoFisher |
| HIF-1$\alpha^{\#}$ | PE | Mouse / IgG1, kappa | Mgc3 | ThermoFisher |
| CD66b* | PE-Cyanine7 | Mouse / IgM, kappa | G10F5 | ThermoFisher |
| MPO$^{\#}$ | PE | Mouse / IgG1 | MPO455-8E6 | ThermoFisher |
| NE$^{\#}$ | Alexafluor 405 | Mouse / IgG1 | 950317R | biotechne |
| CD18* | PE | Mouse / IgG1 | MEM-48 | ThermoFisher |
| CD41a* | Super Bright 436 | Mouse / IgG1, kappa | HIP8 | ThermoFisher |

#### II. Supplementary Figures

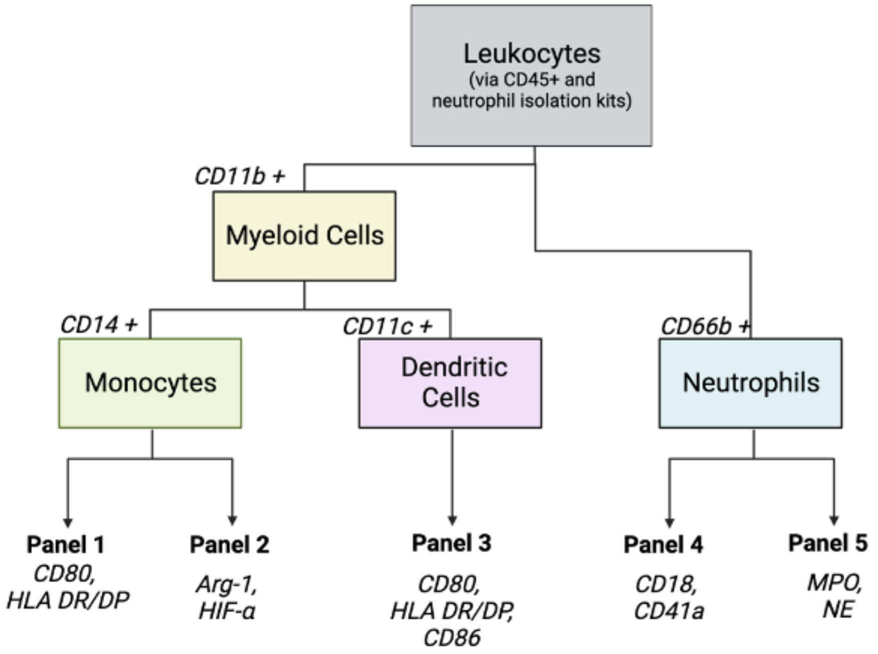


**Figure S1. Panel organization for distinguishing and phenotyping monocytes, dendritic cells, and neutrophils.**

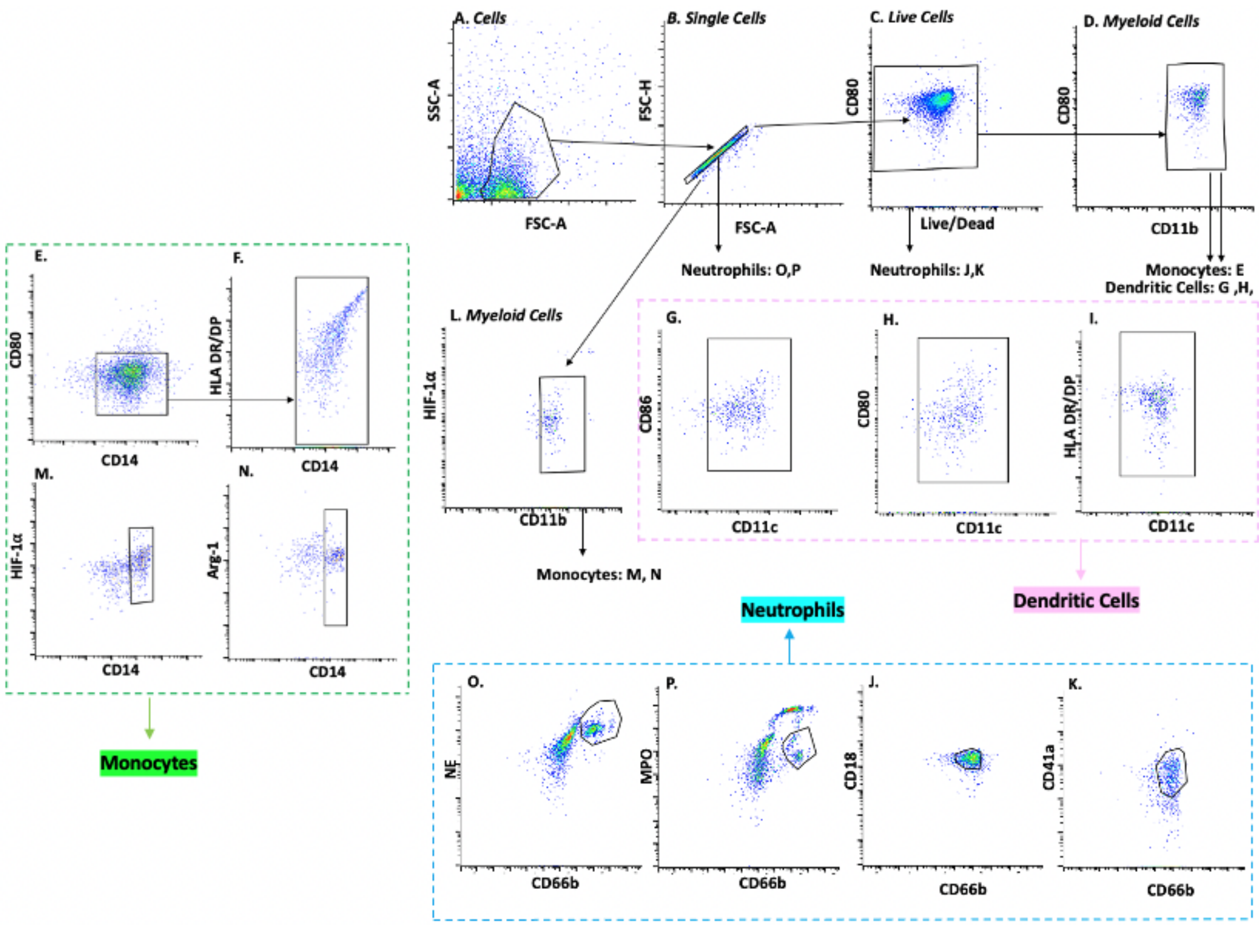


**Figure S2. Immune cell gating.** Hierarchical gates were drawn for (A) cells, (B) single cells, (C) live cells, (D) myeloid cells, (E-F) extracellular markers on monocytes, and (G-I) extracellular markers on dendritic cells. Subplots were drawn from (C) to identify (J-K) extracellular markers on neutrophils. To identify intracellular markers in monocytes, which are inherently non-viable due to the fixation / permeabilization protocol, a subplot was drawn from (B) to identify (L) myeloid cells and (M-N) intracellular markers in monocytes. Subplots were drawn from (B) to identify (O-P) intracellular markers in neutrophils. In the *Main Text,* Figure 3 displays the relative fold-change in median fluorescence intensity for (O) NE, (P) MPO, (J) CD18, and (K) CD41a (blue dotted lines). Figure 4 displays the relative change in median fluorescence intensity for (E) HLA DR/DP, (F) CD80, (M) HIF-1$\alpha$, and (N) Arg-1 (green dotted lines). Figure 5 displays the relative fold-change in median fluorescence intensity for (G) CD86, (H) CD80, and (I) HLA DR/DP (pink dotted lines).

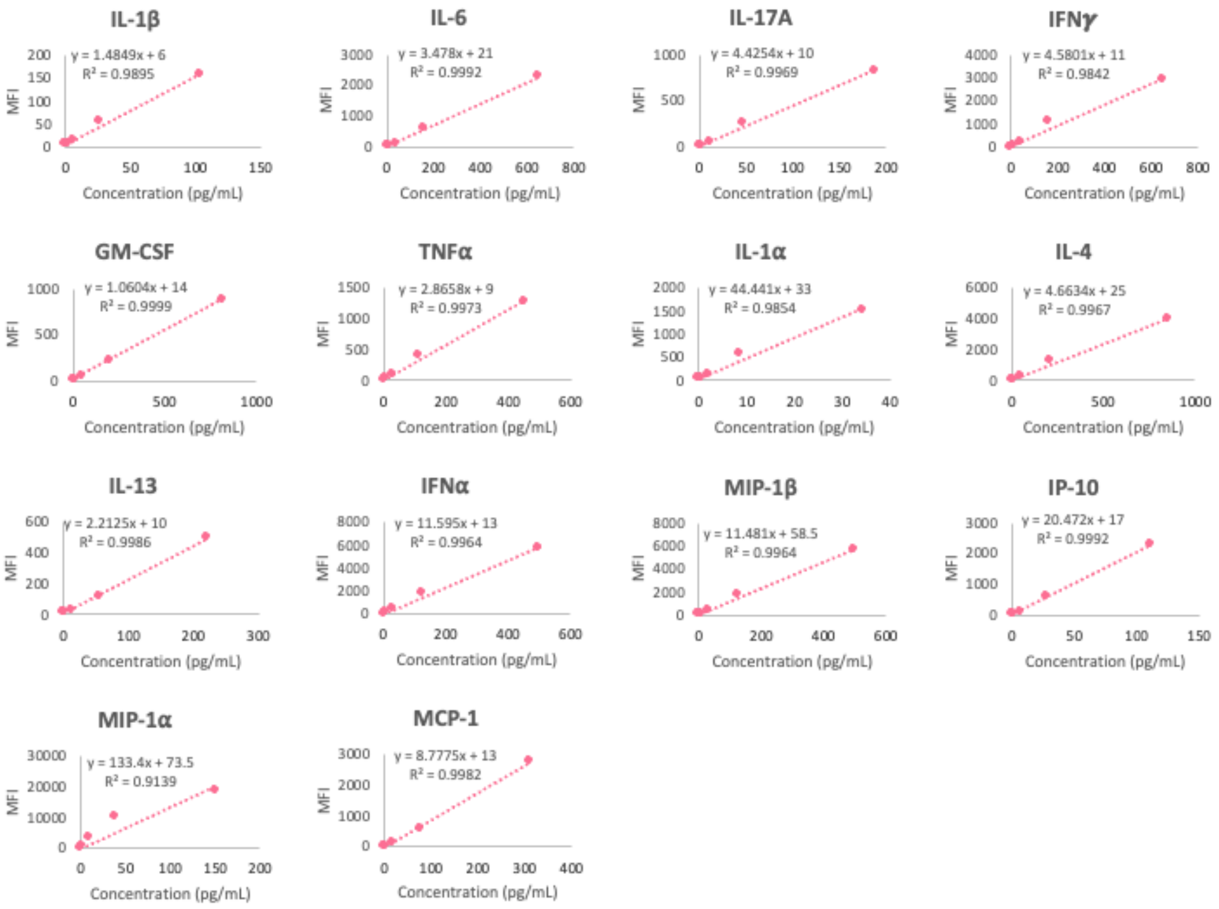


**Figure S3. Standard curves from multiplexed ELISA.** Eight different concentrations were used for each of the fourteen cytokines and chemokines shown in Figure 6 of the *Main Test* to determine standard curves by fitting a linear regression to the data points within the linear regions.
